# Functional Characterization of PCWDEs *VdCut1* and *VdPL16* Reveals Their Critical Roles in *Verticillium dahliae* Pathogenicity

**DOI:** 10.1101/2025.08.22.671736

**Authors:** Gaijie Liu, Jianwei Cao, Xingxing Liu, Jingyi Ye, Yulong Zhang, Asigul Ismayil, Aiying Wang

**Affiliations:** College of Life Sciences, Shihezi University, Shihezi, China; Xinjiang Production and Construction Corps, Key Laboratory of Oasis Town and Mountain-basin System Ecology, Shihezi, China

**Keywords:** *Verticillium dahliae*, PCWDEs, pathogenicity, *VdCut1*, *VdPL16*

## Abstract

Verticillium wilt, a soil-borne fungal disease caused by *Verticillium dahliae*, is a major threat to agricultural production. The plant cell wall serves as the primary barrier against the invasion of fungal pathogens. To facilitate successful infection, many pathogens encode plant cell wall-degrading enzymes (PCWDEs) to degrade cell wall components. In this study, we conducted a genome-wide analysis of PCWDE gene families in two *V.dahliae* strains, VdSHZ-4 and VdSHZ-9, with different pathogenicity. VdSHZ-4 and VdSHZ-9 genomes were found to contain 415 and 396 PCWDE genes, respectively. We successfully isolated two key enzymes from *V. dahliae*: the cutinase *VdCut1* and the pectin lyase *VdPL16*.

To investigate the functions of these two genes, we constructed *VdCut1* and *VdPL16* deletion and complementation mutants and examined their roles in hyphal growth, conidiation, and pathogenicity toward cotton Verticillium wilt. Results showed that deletion of *VdCut1* or *VdPL16* suppressed colony growth and reduced conidial production, and it also impaired the utilization of sucrose and galactose and markedly increased sensitivity to abiotic stresses such as sodium dodecyl sulfate (SDS) and sorbitol. The *VdCut1* deletion mutant completely lost the ability to penetrate cellophane membranes, whereas the *VdPL16* deletion mutant exhibited significantly reduced penetration capacity. Both deletions markedly attenuated virulence, with the loss of *VdCut1* having the more pronounced effect. In planta colonization assays revealed that the *VdCut1* mutant accumulated much less biomass in host tissues than the wild-type strain and the *VdPL16* mutant. These findings demonstrate that *VdCut1* and *VdPL16* play critical roles in *V.dahliae* pathogenicity by regulating hyphal growth and conidiation. The results deepen our understanding of the molecular mechanisms by which this pathogen degrades plant cell walls and provide a theoretical basis for antifungal strategies that target plant cell wall-degrading enzymes (PCWDEs).

**Author summary:** The plant cell wall is the first line of defense against fungal invasion, and *Verticillium dahliae* can secrete plant cell wall-degrading enzymes (PCWDEs) to break this barrier—which is a key step for its successful infection. However, the specific PCWDEs that drive the pathogenicity of *V. dahliae* and their mechanisms of action remain incompletely understood. Here, we conducted a comparative analysis of carbohydrate-active enzymes (CAZymes) in the genomes of two *V. dahliae* strains— (same cotton field origin, high pathogenicity VdSHZ-9 and weak pathogenicity VdSHZ-4). We then focused on key PCWDE genes: *VdCut1* (encoding a cutinase) and *VdPL16* (encoding a pectin lyase), which showed differential pathogenicity associations between the two strains. Using gene deletion mutants and complementation strains, We fund that *VdCut1* and *VdPL16* support *V. dahliae’s* basic biological functions and hyphal penetration, and that the deletion of these genes significantly reduces the pathogenicity of *V. dahliae*. These findings provide a basis for future research on the mechanism underlying the pathogenic variation of *V. dahliae* and identify the genes encoding PCWDEs as potential targets for developing control strategies against cotton Verticillium wilt.

## 1. Introduction

*Verticillium dahliae*, a devastating plant pathogen, induces vascular wilt in diverse crops, significantly impairing agricultural productivity and causing substantial economic losses worldwide ^[1,2]^. Its persistence in soil is enabled by hyphae, conidia, and microsclerotia—specialized survival structures. Among these, microsclerotia are particularly resilient, enduring harsh environmental conditions and remaining viable in soil for over a decade^[3–5]^. Microsclerotia germinate into infectious mycelia, which penetrate host roots through appressoria and penetration pegs, leading to xylem blockage and wilting^[6,7]^. The pathogen’s complex mechanisms make control challenging, underscoring the need to identify key pathogenic genes and molecular pathways. Therefore, identifying key pathogenic genes and elucidating the underlying molecular mechanisms are essential for developing integrated approaches to combat Verticillium wilt^[8]^.

When invading plant cell walls, *V. dahliae* secretes plant cell wall-degrading enzymes (PCWDEs), key virulence factors that degrade pectin, cellulose, and other cell wall components^[9–11]^. This degradation facilitates fungal growth, reproduction, and infection^[12]^. PCWDEs are categorized into pectinases, cellulases, hemicellulases, and cutinases based on their targeted polysaccharide components in the plant cell wall^[13]^. Notably, pathogen pathogenicity often correlates with PCWDE activity; higher activity typically indicates greater pathogenicity^[14]^. Currently, Studies have confirmed that a variety of plant cell wall-degrading enzymes possess pathogenic functions. For example, after knocking out sucrose non-fermenting protein 1 (*VdSNF1*), which is responsible for regulating catabolite repression, the expression levels of a large number of PCWDE genes decrease significantly, and the pathogenic ability of the fungus to plants is also obviously weakened^[15]^. Similarly, compared with the wild-type strain, the knockout mutant of the glucosyltransferase homologous gene VDAG_02071 causes significantly reduced disease symptoms in *Nicotiana benthamiana*^[16]^; When the *VdSSP1* gene is knocked out, the mutant strain shows restricted growth on media with pectin or starch as the sole carbon source, and its pathogenicity to cotton is significantly reduced^[17]^. These research results indicate that *V. dahliae* can infect host plants through a variety of cell wall-degrading enzymes .

All plant-pathogenic fungal PCWDEs belong to the CAZyme superfamily, which is divided into glycoside hydrolases (GHs), carbohydrate esterases (CEs), auxiliary activities (AAs), glycosyl transferases (GTs), polysaccharide lyases (PLs), and carbohydrate-binding modules (CBMs)^[18,19]^. The cuticle covers all aerial parts of plants and serves as the first physical barrier against pathogen entry and infection. It consists of a hydrophobic cutin network formed by esterified hydroxyl and epoxy fatty acid derivatives, which is covered with and partially mixed with wax, acting as a strong barrier to penetration^[20,21]^. To get past the primary obstacle when invading plants, pathogens usually secrete cutinases, extracellular Ser esterases that hydrolyze cutin, which break down the cuticle and help them penetrate^[22–24]^. Comparative analyses of the *V. dahliae* reference genome (strain *VdLs.17*) revealed that it encodes at least fourteen CE5 cutinases, indicating a potentially significant role for these enzymes in pathogenesis^[16]^. Although the involvement of cutinases in the immunomodulatory effects of *V. dahliae* has been investigated, the contribution of CE5 proteins to hyphal growth, carbon-metabolism regulation, and stress adaptation remains largely unknown. Pectin is abundant in fruit peels and the plant cell wall, where it maintains wall integrity and intercellular cohesion^[25,26]^. Because this highly branched polysaccharide is structurally complex, its complete degradation requires a diverse set of enzymes. Consequently, during plant–microbe interactions, pectin-degrading enzymes secreted by phytopathogens play a pivotal role in breaching the plant cell wall^[27]^. Comparative genomics has shown that *V. dahliae* encodes a larger repertoire of carbohydrate-active enzymes (CAZymes) than foliar pathogens, particularly cellulases and pectinases^[28]^. Given their importance during infection, many fungal genes encoding pectin lyases have been identified and characterized. Expression of microbial pectin lyase genes is commonly up-regulated during host colonization and is required for full virulence.

In *V. dahliae*, pectin lyases *VdPL-3.1* and *VdPL-3.3* contribute to cell-wall degradation and pathogenicity^[29]^. Overexpression of the pectin lyase gene *CcpelA* in *Colletotrichum coccodes* enhances virulence^[30]^. *VdPEL1* facilitates pathogen ingress by degrading the plant cell wall^[31]^. Pectate lyase has an inhibitory effect on the *V. dahliae* strain Vd080; it can destroy the structure of spores and mycelia, thereby inhibiting cotton verticillium wilt^[28]^. The *V. dahliae* genome harbors an extensive set of plant cell wall–degrading enzyme genes, and dissecting their functions is essential for elucidating the molecular basis of *V. dahliae* infection.

In this study, we screened PCWDE-related genes in the *V. dahliae* strains VdSHZ-4 (low virulence) and VdSHZ-9 (high virulence) and further filtered pathogenicity-associated genes using annotation results from the Pathogen-Host Interaction (PHI) database. We then conducted a systematic functional analysis of the cutinase gene *VdCut1* (CE5 family) and the pectin lyase gene *VdPL16* (PL1 family). Knock-out mutants (*ΔVdCut1* and *ΔVdPL16*) and their respective complemented strains were generated to evaluate the impact of these genes on fungal growth, carbon-source utilization, abiotic-stress responses, and pathogenicity. Additionally, we conducted signal peptide secretion assays to verify the extracellular activity of these proteins. Our findings clarify the roles of *VdCut1* and *VdPL16* in the growth and pathogenic processes of *V. dahliae*, providing new insights into the molecular mechanisms underlying fungal virulence. This study enhances our understanding of how *V. dahliae* degrades plant cell walls and establishes infection, offering potential targets for developing antifungal strategies.

## 2. Materials and Methods

### 2.1 Plasmids, Fungal Strains and Plant Materials

The *V.dahliae* strains VdSHZ-4 (low virulence) and VdSHZ-9 (high virulence) were isolated from Shihezi, Xinjiang, and their pathogenicity was characterized by the Key Laboratory of Agricultural Biotechnology at Shihezi University. The plasmid pNeo-olic, used for gene complementation, was kindly provided by Prof. Huishan Guo from the Institute of Microbiology, Chinese Academy of Sciences. The cotton cultivar Xinluzao 7 was provided by the Shihezi Academy of Agricultural Sciences, Xinjiang.

### 2.2 Cultivation of *V.dahliae* and Plant Growth Conditions

The strains and their mutants were cultured in liquid Potato Dextrose Agar (PDA) medium at 25°C with shaking at 160 rpm for 7 days^[32]^. The *Agrobacterium tumefaciens* strain GV3101 used in this study was cultured in LB liquid medium containing appropriate antibiotics^[1]^. Cotton seedlings of the variety Xinlu Early No. 7 were grown in a light-controlled greenhouse at 25°C with a 16 h/8 h photoperiod^[33]^.

### 2.3 Identification and Bioinformatics Analysis of PCWDE Genes in *V.dahliae*

#### 2.3.1 Identification of PCWDEs Genes in *V.dahliae*

Based on the genomic sequencing results of VdSHZ-4 and VdSHZ-9 (sequenced on the Illumina NovaSeq 6000 platform using the PE150 sequencing strategy, with a sequencing depth of ≥100X and Contig N50 values of 375,757 bp and 186,711 bp, respectively, S1 Fig), Assembly assessment is shown in S2 Fig. the CAZymes genes in the genomes of VdSHZ-4 and VdSHZ-9 were annotated and classified using the CAZy database (http://www.cazy.org/(accessed on 18 April 2024)), in combination with the dbCAN2 toolkit (v2.0.11), HMMER (E - value ≤ 1×10⁻¹⁵), and DIAMOND (E – value ≤ 1×10⁻⁵) ^[34]^. Potential genes involved in plant cell wall degradation were screened based on the descriptions of degraded substrates in the CAZymes gene annotation information. Meanwhile, a cross-comparison was made with the pathogenicity-related genes reported in the PHI-base database (v4.13, http://www.phi-base.org). The protein sequences of *VdCut1* and *VdPL16* were downloaded from the NCBI website (https://www.ncbi.nlm.nih.gov/). The classical signal peptide structures of PCWDEs genes were predicted using the SignalP 5.0 website (https://services.healthtech.dtu.dk/services/SignalP-5.0/). Non-classical signal peptides were predicted using the SecretomeP 2.0 website (https://services.healthtech.dtu.dk/services/SecretomeP-2.0/). Subcellular localization predictions were made using the WoLF PSORT II tool (https://www.genscript.com/wolf-psort.html). The conserved domains of the *VdCut1* and *VdPL16* genes were predicted using the SMART database (http://smart.embl.de).

#### 2.3.2 Verification of Signal Peptide Activity

Specific primers were used to amplify the signal peptide region sequences of *VdCut1* and *VdPL16*, along with the encoding sequences of the three adjacent downstream amino acids. These sequences were fused with the secretion-deficient invertase vector pSUC2 to construct recombinant vectors, which were then transformed into the yeast strain YTK12^[35]^. The positive yeast strains were streaked on CMD-W (tryptophan-deficient medium) and YPRAA (containing 2% raffinose) plates. The YTK12 yeast strain and the yeast strain containing the empty pSUC2 vector were used as negative controls, while the YTK12 yeast strain harboring the pSUC2::Avr1b^SP^ plasmid served as the positive control. Plates were inverted and cultured at 30°C for 3 days, followed by observation of growth and photography. Additionally, the sucrose invertase activity of the signal peptides (SPs) was detected by the reduction of 2,3,5-triphenyltetrazolium chloride (TTC) to the insoluble red product 1,3,5-triphenylformazan (TPF). The positive colonies were collected, and the cell pellets were stained with 1% TTC (35°C for 10 min, followed by 5 min at room temperature), and the color change of TTC was observed^[36]^. The primers used are listed in S1 Table in the supplementary material.

### 2.4 Analysis of Gene Expression in *V.dahliae* under Different Carbon Source Conditions

Based on Czapek Dox medium (without carbon source), single carbon source solid media were prepared by adding sucrose (30 g/L), galactose (8 g/L), xylose (4.5 g/L), starch (17 g/L), xylan (10 g/L), or pectin (10 g/L) respectively. Each medium was inoculated with 100 µL of a spore suspension (1×10⁷ CFU /mL), spread evenly, and incubated at 25°C for 7 days. Mycelia were collected, RNA was extracted and reverse transcribed into cDNA. Using *β-tubulin* (VDA G_10074) as the reference gene, RT-PCR amplification was performed with specific primers (S1 Table ).

### 2.5 Analysis of Gene Expression Patterns in Cotton Induced by *V.dahliae*

Spore suspension concentration was adjusted to 1×10⁷ CFU/mL. When the cotton seedlings had developed their second true leaf, the root dip method was used to immerse the roots in the *V.dahliae* spore suspension for 15 minutes. The cotton seedlings were then transplanted back into pots and continued to be cultured. Sampling of the roots of inoculated cotton plants was carried out at 3, 12, 24, 48, 72, 96, and 120 hours after inoculation, and the samples were immediately frozen in liquid nitrogen.

Total RNA was extracted from cotton using a plant total RNA extraction kit, and cDNA was synthesized using a reverse transcription kit. Quantitative PCR (qPCR) was performed using ChamQ Universal SYBR qPCR Master Mix (Vazyme Biotech Co., Ltd.) with the primers listed in Table S1. The *VdEF-1α* gene was used as the internal reference. Relative gene expression levels were calculated using the 2^⁻ΔΔCt^ method^[37]^.

### 2.6 Functional Characterization of *VdCut1* and *VdPL16* Genes

#### 2.6.1 Construction of Gene Knockout and Complementation Mutants

Gene knockout and complementary mutants were produced using the *Agrobacterium tumefaciens*-mediated transformation (ATMT)^[38]^. To construct the knockout vectors, the upstream (1125 bp) and downstream fragments of *VdCut1*, as well as the upstream (1219 bp) and downstream flanking fragments of *VdPL16* (1134 bp), were amplified from the genomic DNA of VdSHZ-9 using the primer pairs VdCut1-Up/Down-F/R and VdPL16-Up/Down-F/R, respectively. The Hyg fragment (1.8 kb) was amplified from the pUC-Hyg plasmid using the Hyg-F/R primers (primers are listed in S1 Table). All PCR products were generated using high-fidelity DNA polymerase (Vazyme) and were fused by homologous recombination ^[39]^, and then cloned into the pGKO2-Gateway vector using the Gateway™ BP Clonase™ II kit (Invitrogen). The positive recombinant plasmids were introduced into the GV3101 for fungal transformation^[40]^. Positive transformants were selected on PDA medium containing 50 μg/mL hygromycin and 50 μmol/L 5-fluoro-2’-deoxyuridine (5-F2dU) and verified by PCR using specific primers.

To obtain the complemented strains, fragments containing the promoter, coding region, and terminator sequences of *VdCut1* and *VdPL16* (3,087 and 3,118 bp, respectively) were amplified from the genomic DNA of VdSHZ-9 using the primers pNEO-VdCut1-F/R and pNEO-VdPL16-F/R (S1 Table). The amplified fragments were ligated into the pNeo-olic vector using the ClonExpress II One Step Cloning Kit. The positive recombinant vectors (pNEO-VdCut1/VdPL16) were introduced into the knockout mutants via ATMT and were identified by PCR.

#### 2.6.2 Generation of EGFP-Labeled Strains

Using a plasmid containing the EGFP sequence as a template, a 721 bp EGFP fragment was amplified using primers EGFP-F/R (S1 Table). This fragment was then transformed into VdSHZ-9, as well as the *ΔVdCut1* and *ΔVdPL16* mutant strains, via *Agrobacterium*-mediated transformation. Transformants were verified by PCR. To ensure stable integration and expression of EGFP in *V. dahliae*, positive transformants were subcultured on PDA medium for at least three generations. Fluorescence microscopy with 488 nm excitation was performed to confirm consistent EGFP expression.

#### 2.6.3 Measurement of Colony Growth Rate and Spore Production

To measure the colony growth diameter, wild-type, knockout, and complemented strains of *V. dahliae* were activated on PDA plates. Mycelium plugs (0.5 cm in diameter) were inoculated into 50 mL of Czapek liquid medium and incubated at 25°C with shaking at 160 rpm for 5 days. The center of the PDA solid medium was inoculated with 2 µL of a conidial suspension at a concentration of 1×10⁷ CFU/mL, and the inoculated medium was incubated at 25°C. Colony diameters were measured and recorded at 2, 6, 10, and 14 days post-inoculation. We conducted three replicates for each strain. To determine the spore production of the strains, mycelial plugs (0.5 cm in diameter) from the wild-type, knockout, and complemented mutants of *V. dahliae* were inoculated into 25 mL of Czapek liquid medium. The cultures were incubated at 25°C with shaking at 180 rpm for 3 days. The cultures were then filtered through four layers of gauze, and the spore production of different strains was counted under a microscope using a hemocytometer. Three replicates were set for each strain.

#### 2.6.4 Analysis of Carbon Source Utilization

To assess the carbon source utilization capacity of the strains, mycelial plugs from the wild-type strain VdSHZ-9, knockout strains, and complemented strains were taken from activated petri dishes using a cork borer. These plugs were inoculated into the center of Czapek solid medium (without sucrose) containing sucrose (30 g/L), pectin (10 g/L), starch (17 g/L), cellulose (10 g/L), and galactose (10 g/L). After incubation at 25°C for 14 days, the colony diameters were measured^[41]^. Each treatment was repeated three times.

#### 2.6.5 Assessment of Stress Response Capabilities

Mycelial plugs (0.5 cm in diameter) from the wild-type strain VdSHZ-9, knockout strains, and complemented strains were taken from activated petri dishes using a cork borer. These plugs were inoculated into the center of pre-prepared PDA solid medium containing 1 M sorbitol, 1 M NaCl, 1 M Kcl, 0.02% Congo Red (CR), and 0.002% SDS^[42]^. After incubation at 25°C for 20 days, the colony diameters were measured. Each treatment was repeated three times.

### 2.7 Pathogenicity Assay of *V.dahliae*

#### 2.7.1 Assessment of Penetration Ability

A 3 µL aliquot of spore suspension (1×10⁷ CFU/mL) was inoculated onto a PDA plate covered with sterile cellophane. After drying, the plate was incubated at 25°C for 3 days in a constant-temperature incubator, and the growth of the colonies was observed. Subsequently, the cellophane with hyphae was removed using sterile tweezers, and the plate was further incubated at 25°C for an additional 4 days. Mycelial growth was then observed and photographed^[43]^. Each treatment was replicated three times.

#### 2.7.2 Analysis of Mycelial Colonization in Cotton Vascular Bundles

Cotton plants were infected using the root-dipping method. After 21 days, 2-3 cm segments were excised from the basal stems, thoroughly rinsed with sterile distilled water, and longitudinally sectioned into 0.5–1 mm slice using a sterilized blade. The sections were stained in 50 μM propidium iodide (PI) prepared in phosphate-buffered saline (PBS, pH 7.4) for 30 minutes in the dark, followed by three rinses with fresh PBS. After mounting, hyphal colonization within the cotton vascular bundles was observed using a Nikon AXR laser scanning confocal microscope^[44]^.

#### 2.7.3 Pathogenicity Assay

The root-dipping inoculation method was used to inoculate cotton plants with a conidial suspension (1×10⁷ conidia/mL) of the wild-type, deletion mutant, and complementary strains. On the 21st day after inoculation, the cotton disease index (DI) was statistically analyzed using the five-grade classification method^[45]^. The formula is: DI = ∑[(the number of diseased plants at each grade × the representative value of the disease grade) / (the total number of plants × 4)] × 100%^[46]^. At the same time, the degree of browning inside the stem was checked and recorded. Moreover, the cotton stems 21 days after inoculation were surface-sterilized, then cut into segments approximately 1 cm in length, and neatly placed on PDA medium supplemented with antibiotics. After incubation at 25°C for 5 days, the fungal mycelia produced by the stem segments were observed and photographed^[47]^. The experiment for each strain was repeated three times, with four pots for each strain, and three cotton seedlings planted in each pot.

## 3. Results

### 3.1 Comparative Analysis of CAZy Gene Families in VdSHZ-9 and VdSHZ-4

Carbohydrate-active enzymes (CAZymes) are a class of proteins capable of degrading and catalyzing polysaccharides. In the genome of *V. dahliae* strain VdSHZ-9, we identified 645 putative genes encoding CAZymes, including 298 glycoside hydrolases (GHs), 103 glycosyltransferases (GTs), 53 carbohydrate esterases (CEs), 73 auxiliary activities enzymes (AAs), 31 polysaccharide lyases (PLs), and 87 carbohydrate-binding modules (CBMs). In contrast, the genome of VdSHZ-4 contained 638 CAZy genes, comprising 296 GHs, 100 GTs, 56 CEs, 70 AAs, 33 PLs, and 83 CBMs (Figure 1A, Supplementary Table S2). A total of 114 CAZy subfamilies were shared between VdSHZ-9 and VdSHZ-4 (Fig 1B).

**Fig 1.**
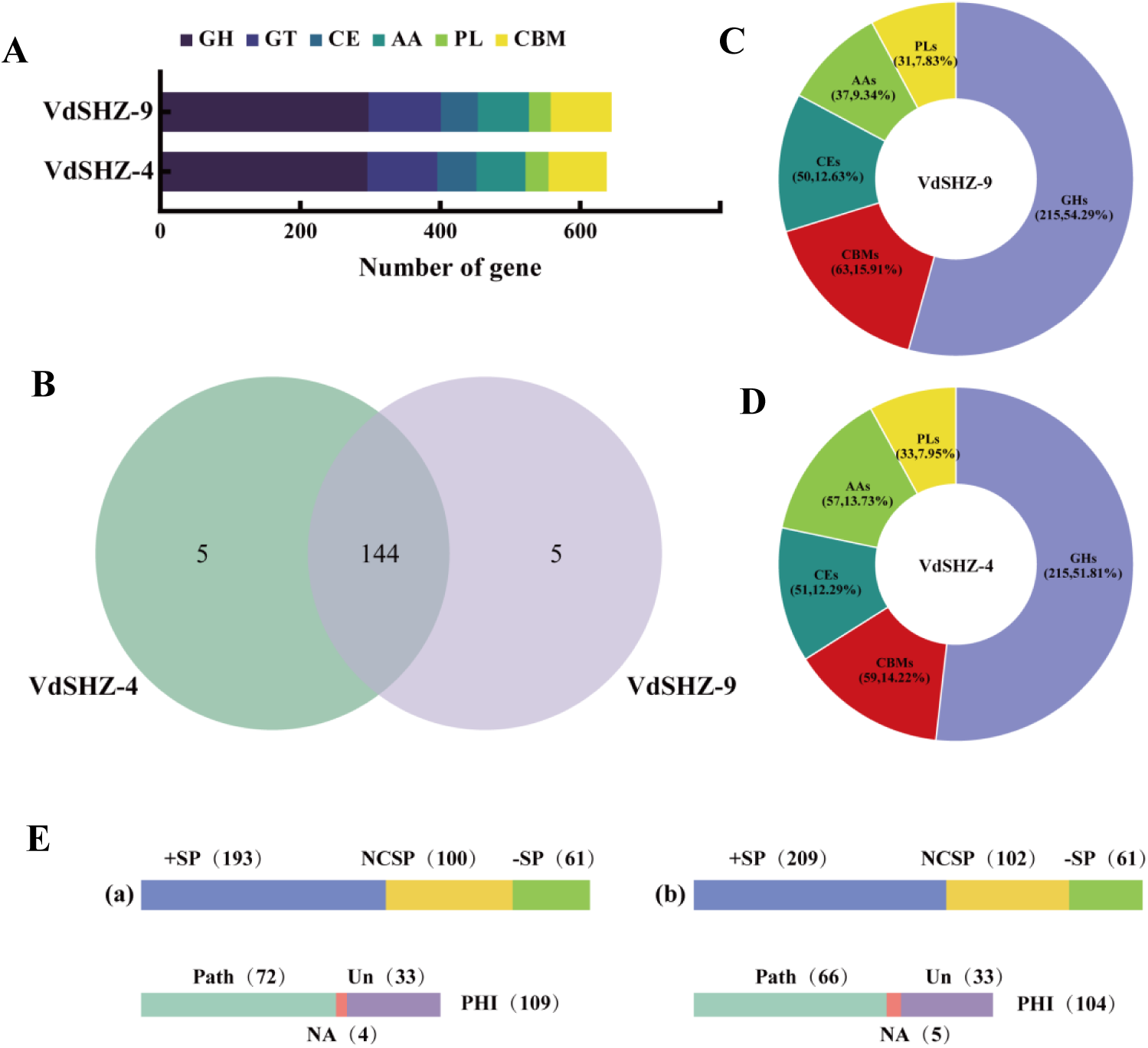
Annotation of Plant Cell Wall Degrading Enzymes (PCWDEs) in the Genomes of VdSHZ-4 and VdSHZ-9. **(A)** Composition of CAZymes families in VdSHZ-4 and VdSHZ-9. Glycoside Hydrolases (GH); Carbohydrate Esterases (CE); Auxiliary Activities (AA); Glycosyl transferases (GT); Polysaccharide Lyases (PL); Carbohydrate-Binding Modules (CBM)^[48]^. **(B)** Subfamily distribution of CAZymes in *V. dahliae*. **(C)** Distribution of PCWDE genes in VdSHZ-9. **(D)** Distribution of PCWDE genes in VdSHZ-4. **(E)** Signal peptide prediction and PHI protein annotation in SHZ-9 (a) and SHZ-4 (b). Upper panel: Signal peptide prediction of PCWDE genes, including classical signal peptides (+SP), non-classically secreted proteins (NCSP), and proteins without signal peptides (-SP). Lower panel: PHI protein annotation of PCWDEs, categorized as pathogenicity-related (Path), no homologous proteins (NA), or non-pathogenicity-affecting (Un).

### 3.2 Identification of PCWDE Genes and Pathogenicity Association Analysis in VdSHZ-9 and VdSHZ-4

Based on the substrate specificity of CAZyme genes, the genome of strain VdSHZ-9 is predicted to encode 396 putative PCWDEs (Fig 1C, S3 Table), whereas the genome of strain VdSHZ-4 encodes 415 putative PCWDEs (Fig 1D, S4 Table). These PCWDEs can be classified into five functional categories, among which the GH family represents the largest group, accounting for more than 50% of all PCWDEs in both strains.

For VdSHZ-9, signal-peptide analysis of the 354 PCWDEs revealed that 58 lack a recognizable signal peptide, 193 possess a canonical signal peptide, and 100 are predicted to be non-classically secreted proteins. Additionally, 109 of these genes were annotated as PHI-base proteins, 72 of which are associated with fungal pathogenicity (Fig 1E-a, S5 Table). Similarly, within the 372 PCWDEs of VdSHZ-4, 209 contain a canonical signal peptide and 102 are predicted as non-classically secreted proteins. Among them, 104 were annotated as PHI-base proteins, with 66 linked to fungal pathogenicity (Fig 1E-b, S6 Table ).

### 3.3 PHI Protein Comparison and Pathogenic Gene Mining in VdSHZ-9 and 4

A total of 109 PHI proteins were identified in VdSHZ-9 and 104 in VdSHZ-4. Comparative analysis revealed that 85 genes exhibited consistent PHI functional annotations in both strains. VdSHZ-9 and VdSHZ-4 harbored 19 and 14 unique PHI genes, respectively. Additionally, five genes displayed inconsistent PHI annotations between the two strains (Fig 2A). Among these five, *A6444* (PHI:2383) and *A0452* (PHI:4619) were linked to the pathogenicity of VdSHZ-9 but showed no effect on VdSHZ-4.

**Fig 2.**
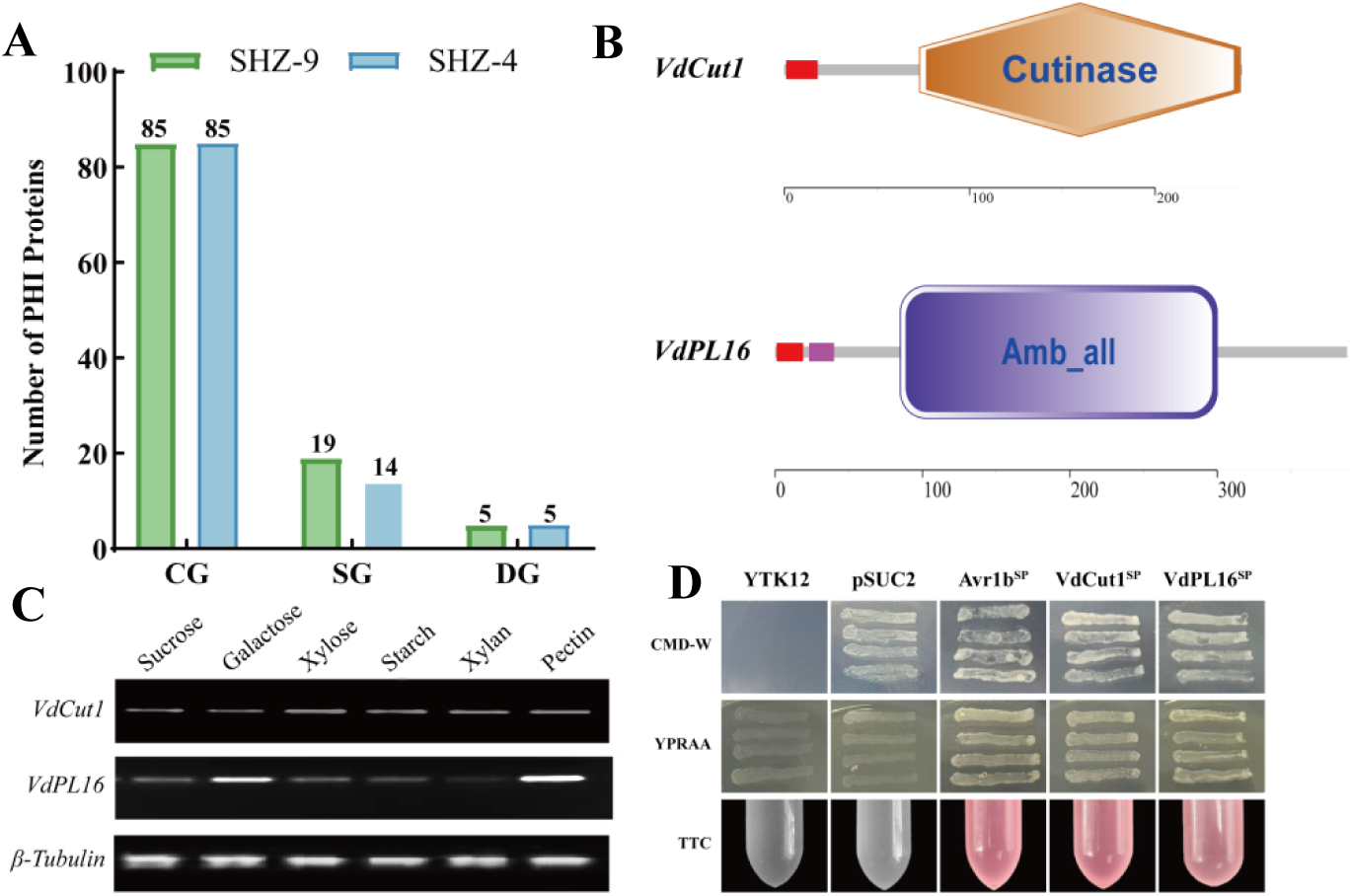
Comparison of PHI Gene differences Between VdSHZ-9 and VdSHZ-4, Functional Domain Prediction, Carbon-Regulated Expression, and Secretion Validation of *VdCut1* and *VdPL16*. **(A)** Comparison of PHI genes between VdSHZ-9 and VdSHZ-4. CG: Common PHI genes; SG: Specific PHI genes; DG: PHI genes with different annotation results. **(B)** Based on the Pfam database, the trehalase domains in *VdCut1* and *VdPL16* were predicted. **(C)** RT - PCR analysis of the gene expression of *VdCut1* and *VdPL16* under different single-carbon-source conditions. **(D)**The functionality of the signal peptide in *VdCut1*and *VdPL16* was verified through the yeast signal trap system.

*A6444* (VDAG_03117), designated *VdCut1*, encodes a cutinase. Its 744-bp full-length cDNA encodes a 247-aa protein with an N-terminal 19-aa signal peptide and an Abhydrolase superfamily domain. *A0452* (VDAG_08154), named *VdPL16*, contains a 1167-bp full-length cDNA encoding a 388-aa protein with a 20-aa N-terminal signal peptide and an Amb_all domain belonging to the PL-6 superfamily (Fig 2B).

In addition, *VdPL16* was highly induced by galactose and pectin, whereas *VdCut1* expression remained stable across all carbon sources tested (Fig 2C). Yeast signal trap assays demonstrated that the signal peptides of *VdCut1* and *VdPL16*, when fused with the invertase gene (lacking a signal peptide in the plasmid pSUC2) in yeast, possessed the ability to mediate invertase secretion and successfully conferred the yeast strain YTK12 with the capacity to grow normally on a medium with raffinose as the sole carbon source (Fig 2D).

### 3.4 *VdCut1* and *VdPL16* are essential for colony growth and conidial formation

To investigate the biological functions of *VdCut1* and *VdPL16*, targeted gene-deletion mutants were generated via homologous recombination. PCR screening identified two independent knockout lines for each gene (*ΔVdCut1-1*, *ΔVdCut1-2* and *ΔVdPL16-1*, *ΔVdPL16-2*). All strains were grown at 25 °C in darkness for 14 d. Compared with the wild-type strain VdSHZ-9, colony morphology of *ΔVdCut1-1/2*, *ΔVdPL16-1/2* and their corresponding complemented strains (*C-ΔVdCut1* and *C-ΔVdPL16*) showed no obvious differences, and all produced dense aerial mycelia (Fig 3A).

**Fig 3.**
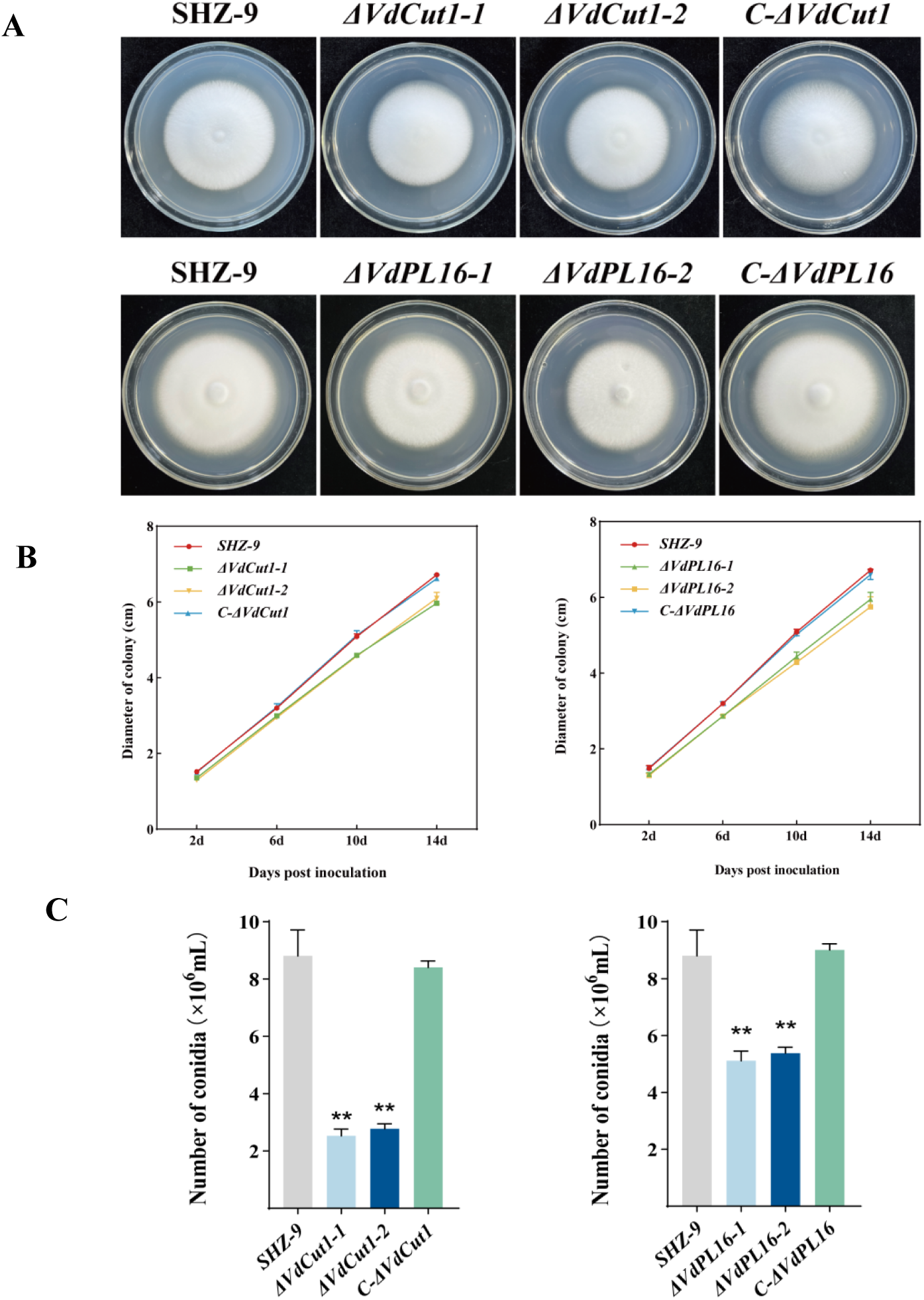
Evaluation of colony morphological and measurement of conidial yields. **(A)** Colony morphology of WT, *ΔVdCut1-1/2*, *ΔVdPL16-1/2*, *C-ΔVdCut1*, and *C-ΔVdPL16* strains in PDA after being cultured at 25 °C for 14 days. **(B)** Colony diameters of different *V. dahliae* strains in PDA at different time points. **(C)** Conidia yields of distinct *V. dahliae* strains following 4 days of cultivation on Czapek medium. Means and standard errors from three biological replicates are shown. Asterisks (**) indicate significant differences (P < 0.01).

Colony diameters measured by the cross method revealed that the growth of *ΔVdCut1-1/2* and *ΔVdPL16-1/2* mutants became increasingly retarded over time relative to the wild type (Fig 3B). In addition, conidiation assays showed that the conidial yields of *ΔVdCut1-1/2* and *ΔVdPL16-1/2* mutants were significantly lower than those of the wild type and the complemented strains (Fig 3C).

### 3.5 Carbon-source utilization by *ΔVdCut1* and *ΔVdPL16* mutants

To assess the capacity of *ΔVdCut1* and *ΔVdPL16* strains to utilize different carbon sources, each strain was inoculated onto Czapek solid media supplemented with the indicated carbon sources and cultured for 14 d (Fig 4A). Colony morphology and radial growth were then compared against the wild-type VdSHZ-9. On sucrose medium, radial growth of *ΔVdCut1-1/2* was significantly reduced and the mycelium appeared sparse compared with VdSHZ-9. By contrast, on galactose medium the *ΔVdCut1-1/2* mutants grew faster than both the wild-type and the complemented strains, suggesting that *VdCut1* negatively regulates galactose utilization in *V.dahliae*.

**Fig 4.**
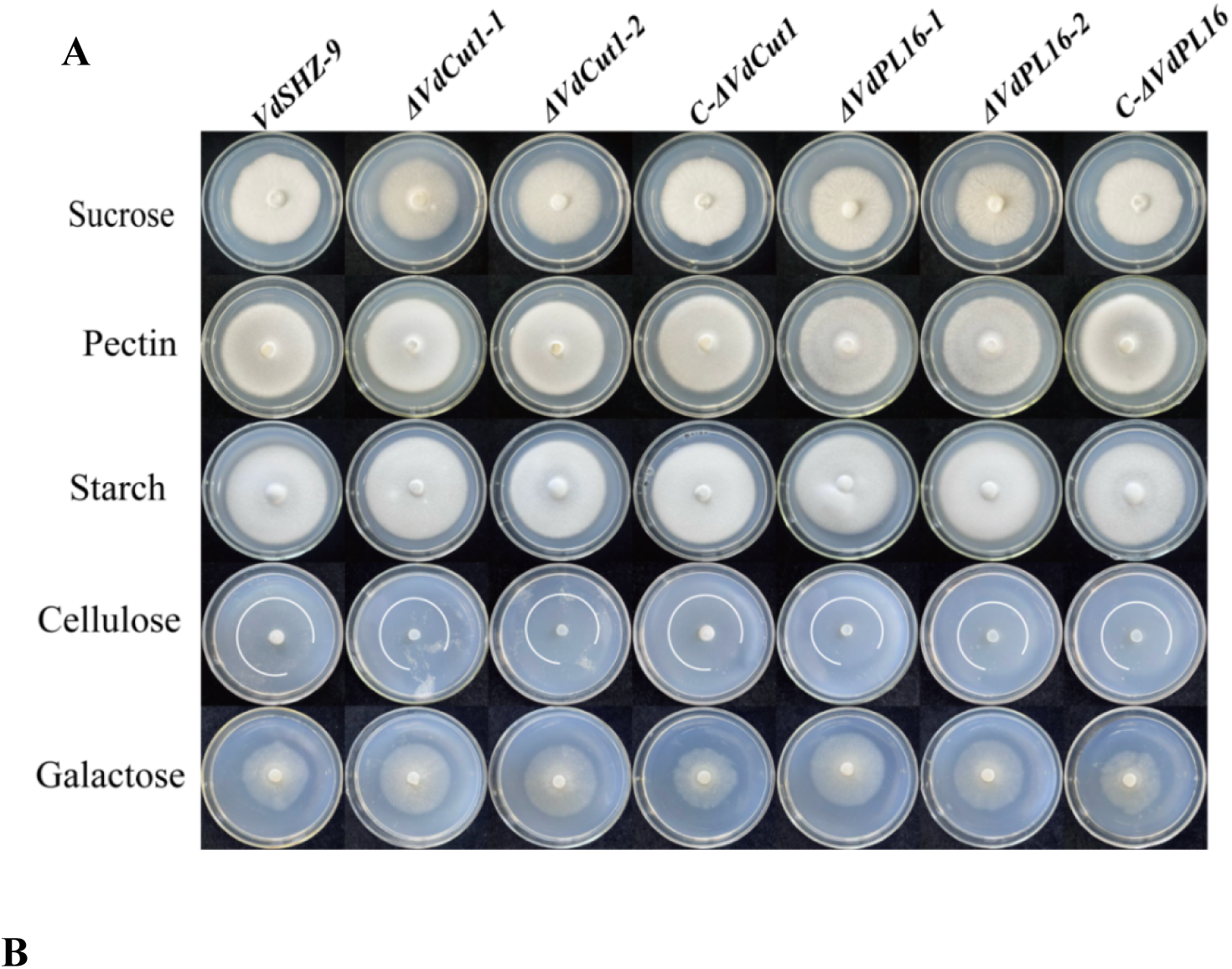
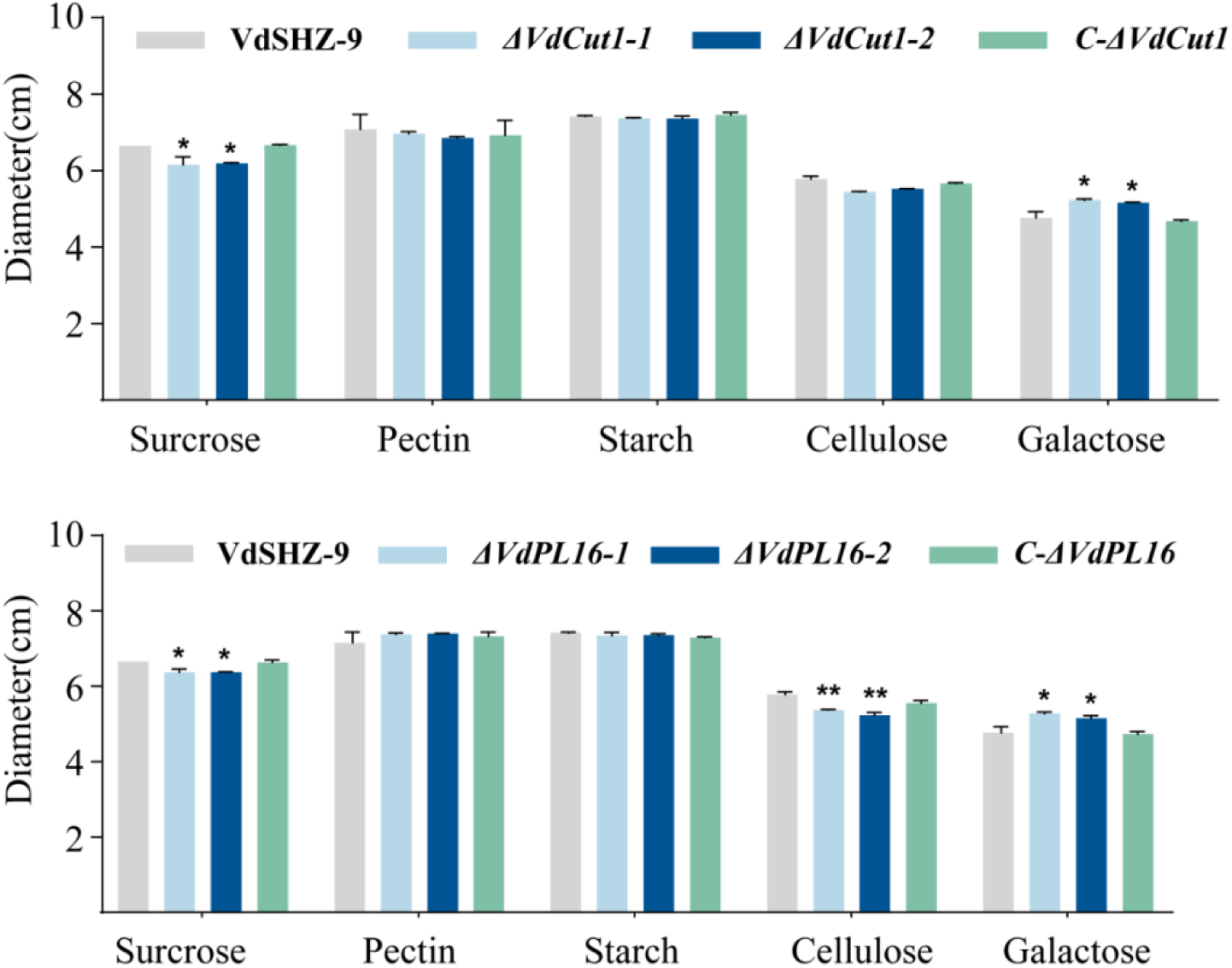
Analysis of the roles of *V. dahliae* strains in the utilization of different carbon sources. **(A)** Growth phenotypes of VdSHZ-9, *ΔVdCut-1/2* and *ΔVdPL16-1/2* grown on media supplemented with various carbon sources. **(B)** The diameters of growth for each tested strain. The mean and standard error calculated from three independent biological replicates are displayed. Asterisks (*) signify significant differences at P < 0.05, (**) represent significant differences (P < 0.01).

The *ΔVdPL16-1/2* mutants exhibited markedly impaired growth on sucrose and cellulose as sole carbon sources. When pectin was supplied, the *ΔVdPL16-1/2* colonies were also notably smaller and less dense (Fig 4B).

Collectively, these results indicate that deletion of either *VdCut1* or *VdPL16* compromises the ability of *V.dahliae* to utilize a broad spectrum of carbon sources.

### 3.6 Stress tolerance of *ΔVdCut1* and *ΔVdPL16* mutants

After 14 d of incubation on different stress media, colony morphology and growth of VdSHZ-9, *ΔVdCut1-1/2* and *ΔVdPL16-1/2* were documented (Fig. 5A). Compared with the wild type, *ΔVdCut1-1/2* exhibited significantly reduced growth on media supplemented with Kcl, sorbitol, Congo red (CR) or SDS; complementation fully restored the wild-type phenotype. *ΔVdPL16-1/2* was likewise impaired under sorbitol, CR and SDS stresses, and growth was almost completely arrested in the presence of sorbitol or CR (Fig 5B).

**Fig 5.**
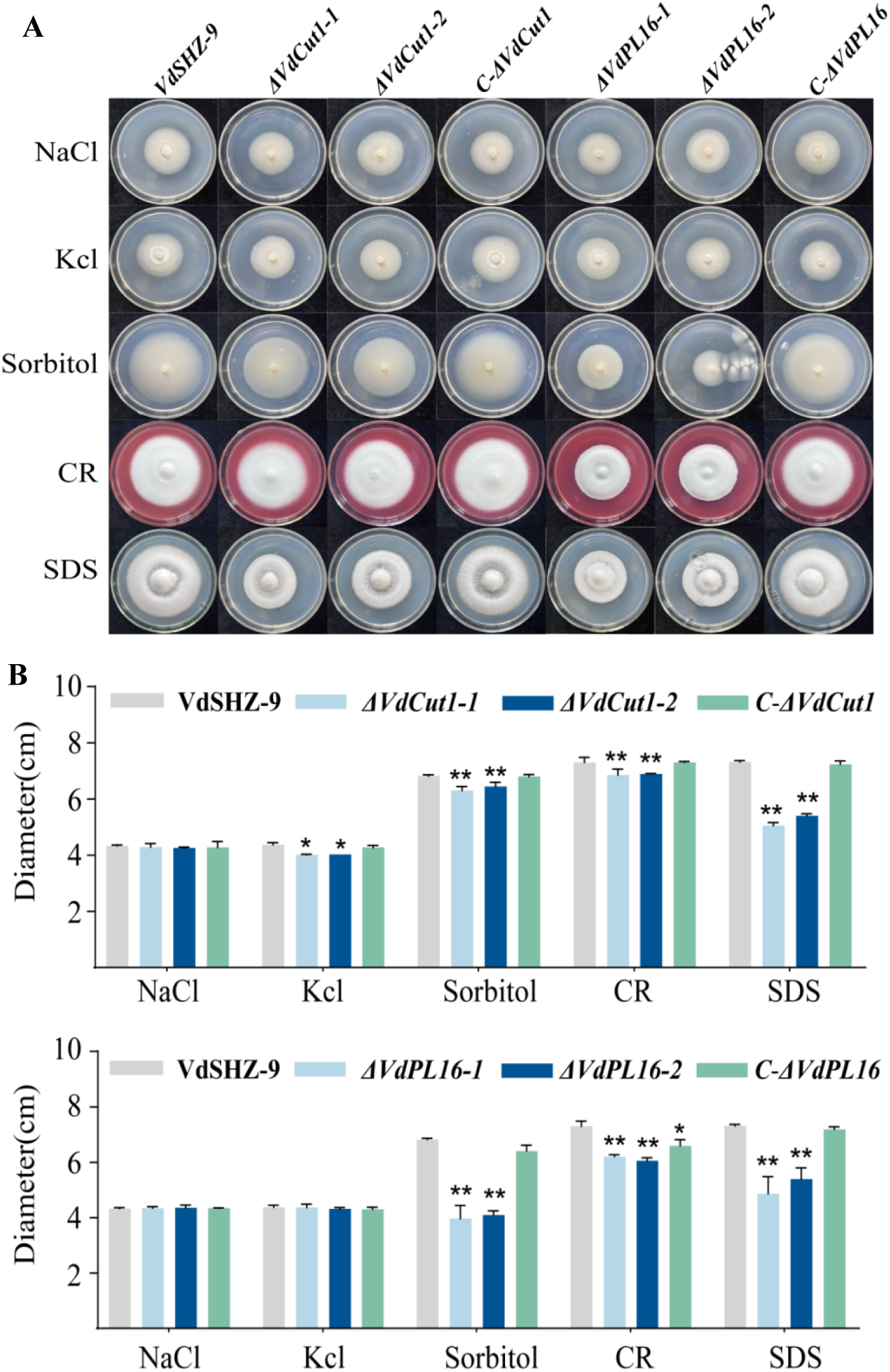
Examination of colony morphology and growth dynamics of *V. dahliae*e strains under different stress Conditions. **(A)** Morphological characteristics of colonies from different *V. dahliae*e strains in multiple stress factors. **(B)** Growth rate of diverse *V. dahliae*e strains under differential stress regimens. Means and standard errors from three biological replicates are shown. Asterisks (*) indicate significant differences (P < 0.05), (**) indicate significant differences (P < 0.01)

Taken together, deletion of either *VdCut1* or *VdPL16* markedly decreases the tolerance of *V.dahliae* to osmotic (Kcl, sorbitol), cell-wall (CR) and membrane (SDS) stresses, indicating that both genes are required for the fungus to withstand high-salinity, hyperosmotic and cell-surface-perturbing conditions.

### 3.7 *VdCut1* and *VdPL16* are crucial for hyphal penetration and host vascular colonization of *V. dahliae*

The contribution of *VdCut1* and *VdPL16* to hyphal penetration was evaluated with a cellophane-membrane assay. Three days after membrane removal, hyphae of the wild-type strain VdSHZ-9 and the complemented strains *C-ΔVdCut1* and *C-ΔVdPL16* readily penetrated the membrane and continued to grow on the underlying PDA medium. In contrast, the *ΔVdCut1-1/2* mutants completely failed to breach the membrane, whereas *ΔVdPL16-1/2* mutants retained partial but significantly reduced penetration capacity (Fig 6).

**Fig 6.**
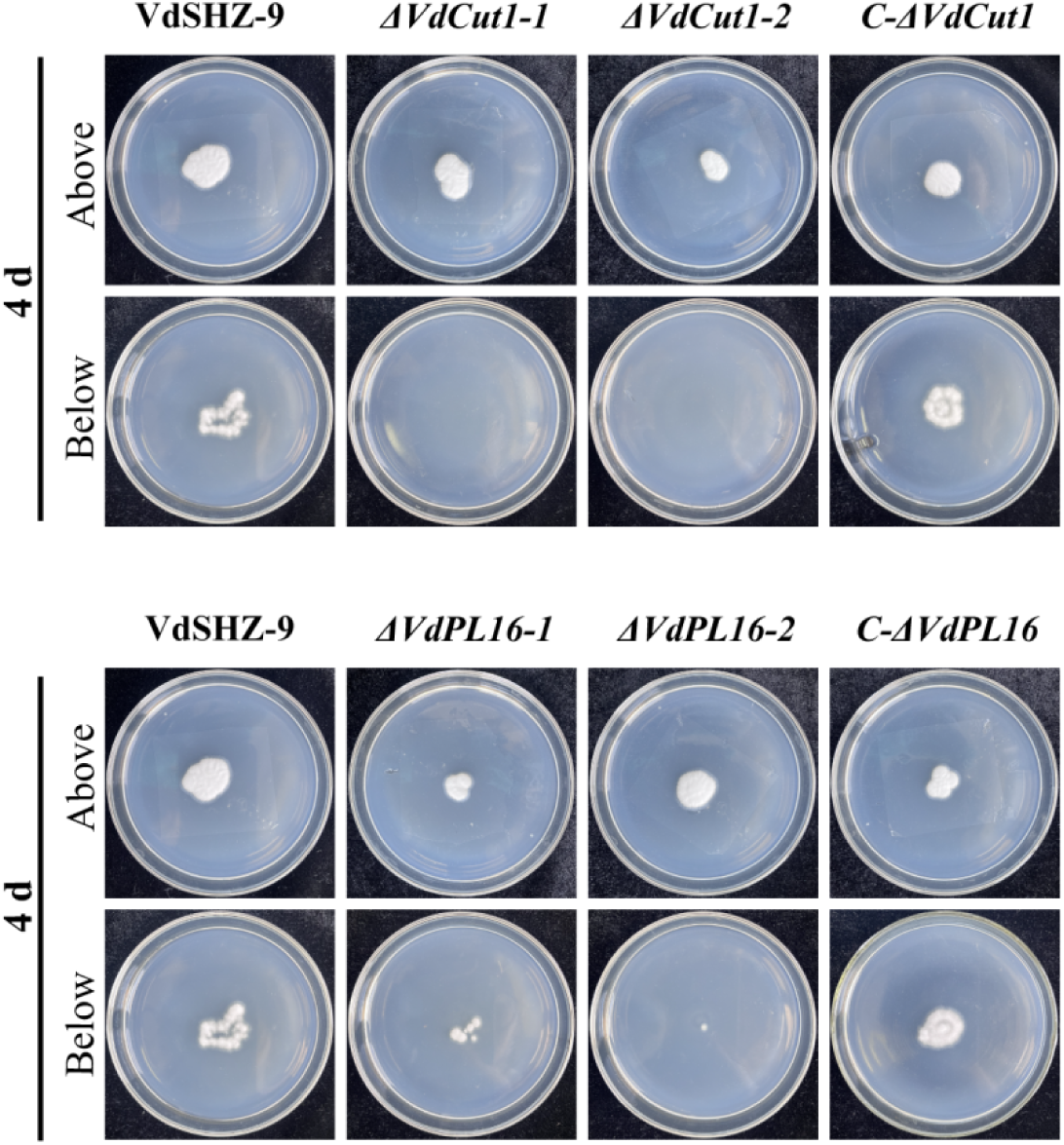
Assessment of the penetration capacity of VdSHZ-9 and mutants on Potato Dextrose Agar (PDA) medium overlaid with cellophane (Above) or without cellophane (Below). These strains were cultured on PDA medium plates overlaid with cellophane at 25 °C for 3 days, and then further cultured for 4 days after the cellophane was removed.

To examine the impact on host colonization, cotton seedlings were inoculated with GFP-expressing strains. At 21 days post-inoculation (dpi), abundant hyphae were observed in the xylem vessels of stems inoculated with VdSHZ-9-EGFP. However, no hyphal signals were detected in plants inoculated with *ΔVdCut1*-EGFP, indicating a complete loss of vascular colonization. Plants inoculated with *ΔVdPL16*-EGFP displayed only sparse hyphal networks within the xylem vessels (Fig 7).

**Fig 7.**
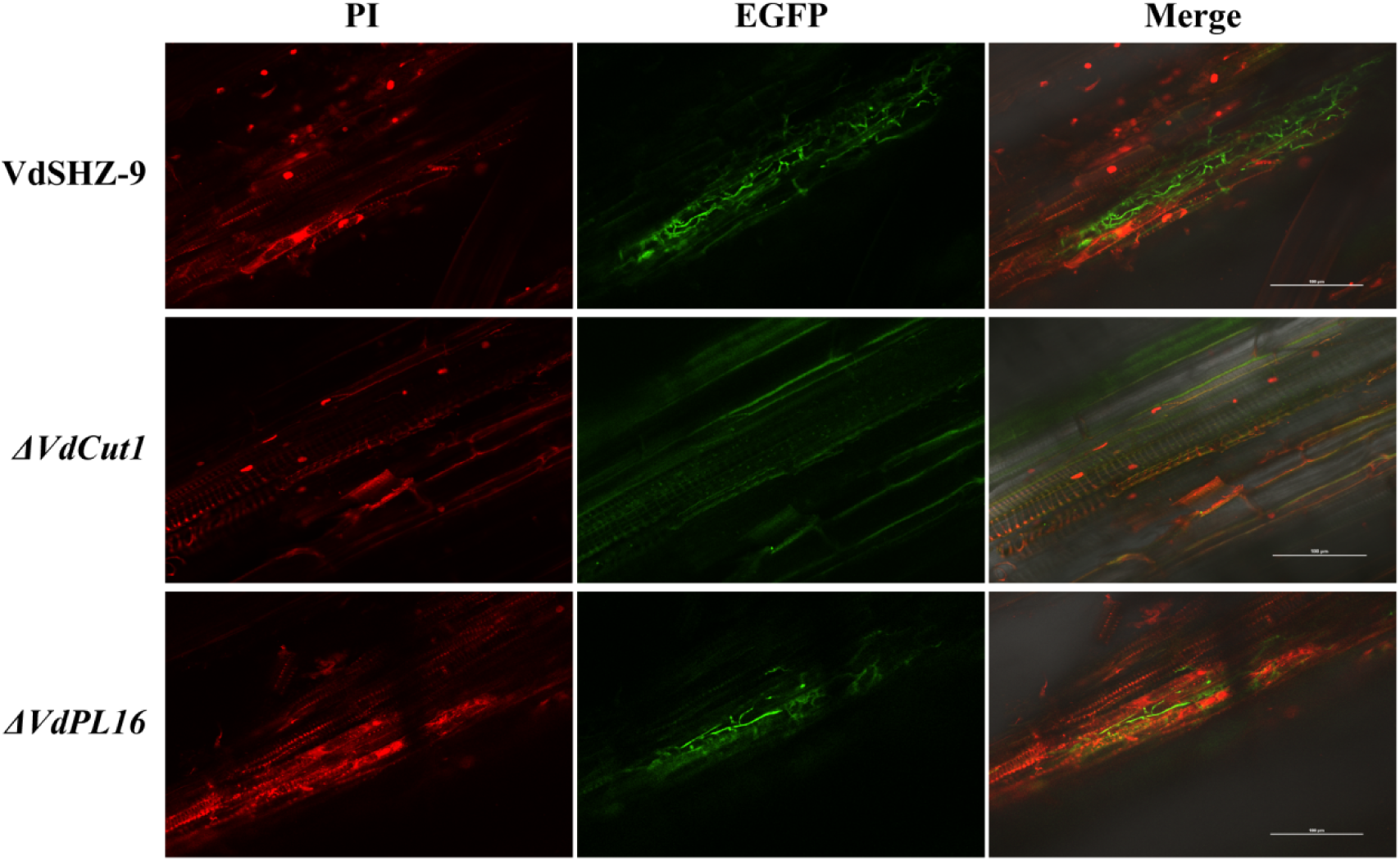
Colonization of different strains in cotton. Longitudinal sections of the stem microtubule tissues of cotton after inoculation with VdSHZ-9-EGFP, *ΔVdCut1*-EGFP, and *ΔVdPL16*-EGFP strains for 21 days. Scale bar, 100 μm.

Taken together, these findings demonstrate that *VdCut1* and *VdPL16* are indispensable for the initial penetration of *V. dahliae* into root epidermal cells and for subsequent systemic colonization and spread within the host vascular system.

### 3.8 Deletion of *VdCut1* or *VdPL16* attenuates the virulence of *V. dahliae*

The distorted hyphal morphology and compromised penetration ability of the knockout mutants suggested that their pathogenicity might be impaired. At 21 days post-inoculation (dpi), mock-inoculated cotton seedlings remained healthy and symptom-free, whereas plants challenged with the wild-type (WT) strain VdSHZ-9 or the complemented strains (*C-ΔVdCut1* and *C-ΔVdPL16)* displayed typical Verticillium wilt symptoms, including wilting, necrosis, and vascular browning (Fig. 9A). In contrast, cotton inoculated with either *ΔVdCut1-1/2* or *ΔVdPL16-1/2* mutants developed only mild wilting and chlorosis. Notably, vascular browning was more pronounced in *ΔVdPL16*-infected plants than in those infected with *ΔVdCut1* (Fig 8A). A detailed disease assessment revealed that cotton seedlings inoculated with the VdSHZ-9 strain predominantly exhibited disease severity grades of 3 and 4, with a disease index (DI) of 90. The complemented mutant strains (*C-ΔVdCut1* and *C-ΔVdPL16*) also caused relatively high disease indices, with values of 88.33 and 85, respectively. In contrast, cotton seedlings inoculated with the knockout mutant strains *(ΔVdCut1-1*, *ΔVdCut1-2*, *ΔVdPL16-1*, and *ΔVdPL16-2*) displayed significantly milder disease symptoms, with disease severity grades mainly ranging from 1 to 2. A small number of cotton plants inoculated with the *ΔVdPL16* strain showed a disease severity grade of 3. The corresponding disease indices for these knockout mutant strains were 30, 31.67, 38.33, and 36.67, respectively (Fig 8B).

**Fig 8.**
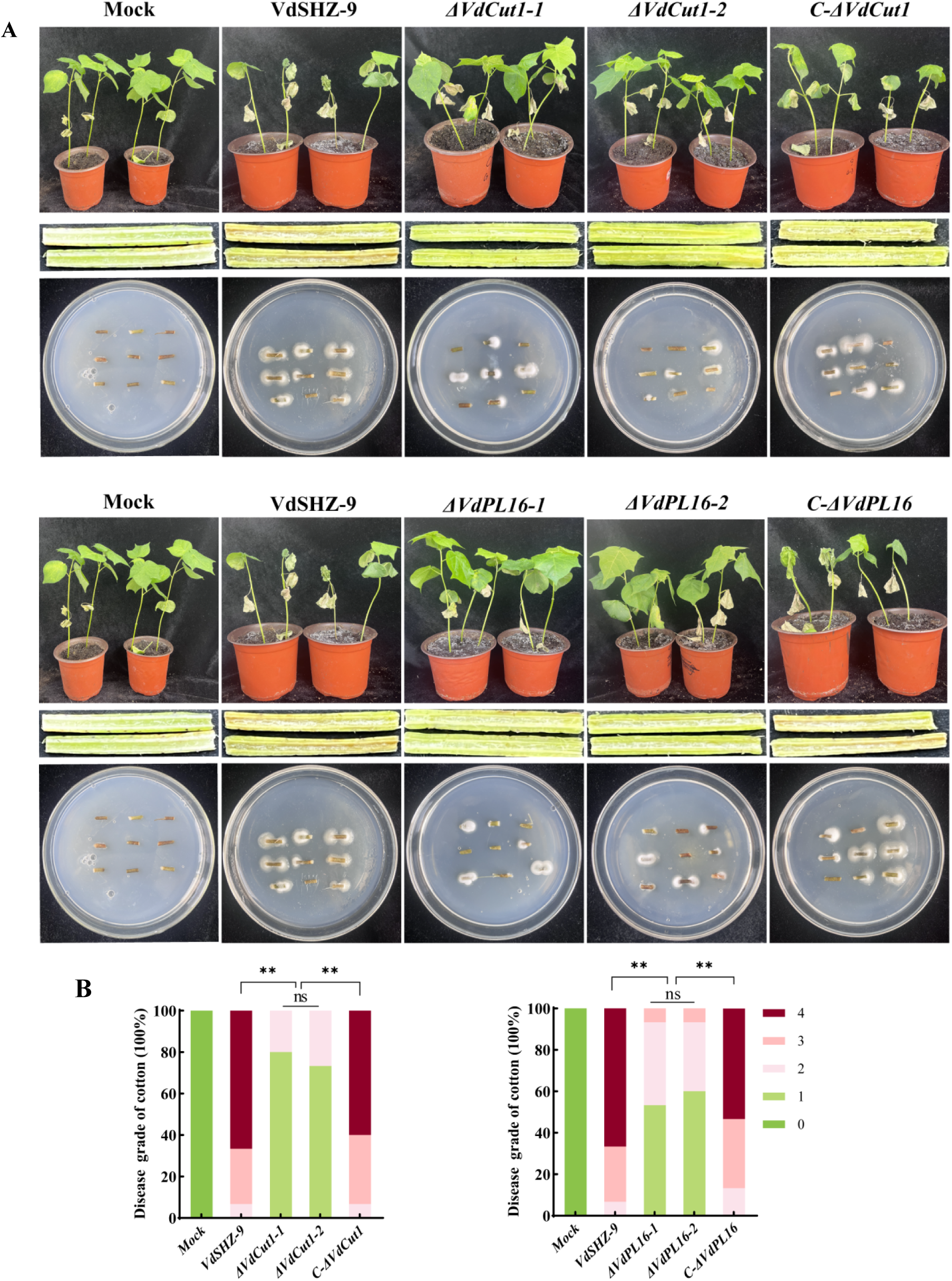
Assessment of the pathogenicity of various strains following infection of cotton. **(A)** Pathogenic phenotypes (upper part), vascular bundle browning (middle part), fungal isolation experiments (lower part) of cotton after inoculation with sterile water (Mock), wild-type strain (VdSHZ-9), knockout strains (*ΔVdCut1-1/2* and *ΔVdPL16-1/2*), and complementary strains (*C-ΔVdCut1* and *C-ΔVdPL16*) for 21 days. **(B)** Disease grades of infected cotton plants at 21 days. The asterisks indicate statistical differences compared with the wild-type and complementary mutant strains. Asterisks (**) indicate significant differences (P < 0.01).

These results provide compelling evidence that the deletion of *VdCut1* and *VdPL16* significantly reduces the pathogenicity of *V.dahliae*. The knockout mutants exhibited a marked decrease in their ability to cause severe disease symptoms in cotton plants, highlighting the crucial roles of these genes in the infection and disease development processes of this devastating pathogen.

## 4. Discussion

Carbohydrate-active enzymes (CAZymes) catalyse the synthesis, modification or breakdown of carbohydrates^[38,49]^. Their repertoire largelydetermines a fungus’s competitiveness, growth and reproduction. Most phytopathogenic fungi secrete plant cell-wall-degrading enzymes (PCWDEs) to breach and penetrate host cell walls^[50,51]^. Comparative genomics has shown that the *V.dahliae* genome encodes more CAZymes than many other fungi^[52]^. Numerous studies have linked PCWDEs to enhanced cell-wall degradation^[53,54]^. In this work, CAZymes and PCWDEs were predicted for the two strains that differ in virulence. VdSHZ-9 and VdSHZ-4 harbour 645 and 638 CAZyme family genes, respectively, together with 396 and 415 PCWDE genes. Genome annotation revealed a high degree of conservation: the two strains share 144 CAZyme families, indicating either a short divergence time or strong stabilizing selection within the species^[52]^. Comparing genomes of strains grown under identical conditions but differing in virulence allows the identification of loci that drive pathogenicity. Minor functional differences can be dissected to reveal the genetic basis of virulence divergence^[55,56]^.

Among PHI-base–annotated genes, the relationship between shared PHI categories and virulence is of particular interest; loss-of-function or down-regulation of most PHI genes attenuates or abolishes pathogenicity^[57]^. PHI annotation of the predicted PCWDEs identified 109 and 104 PHI genes in VdSHZ-9 and VdSHZ-4, respectively, falling into 70 and 67 PHI classes (PHI IDs). Cross-strain comparison showed that most PHI genes were numerically conserved, but five differed in their pathotype annotation. Notably, PHI:2383 has been implicated in stone-fruit wilt: its inactivation markedly reduces virulence. Likewise, PHI:4619 attenuates *Phytophthora capsici* pathogenicity^[58,59]^. These observations suggest that the distinct pathotypes of these PHI genes may underlie the virulence difference between the two strains.

Cotton Verticillium wilt begins with root invasion followed by proliferation in xylem vessels^[60]^. Cutinases play a key role in the interaction between fungi and plant hosts. They not only can degrade plant cuticles or suberin polymers, induce host signal transmission, assist fungal spores in attaching to plant surfaces, and acquire carbon sources during the saprophytic stage, but also participate in the process of host infection, making them an indispensable factor for pathogens to develop virulence^[61–63]^. Studies have found that *VdCUT11* hydrolyses suberin model compounds, the products acting as DAMPs that trigger plant immunity and are strongly up-regulated 1–2 dpi^[64]^. Here, *VdCut1* and *VdPL16* were also markedly up-regulated at 1–2 dpi (S3 Fig), implying a possible functional coupling during early infection. Conidia and hyphae are critical for infection; genes such as *VdPT1* (neutral trehalase), *VdNuo1* (NADH-ubiquinone oxidoreductase) and *VdGAL4* affect radial growth, conidiation and penetration^[1,3,65]^. Deletion of *VdCut1* or *VdPL16* reduced colony expansion and conidial yield without altering aerial hyphal morphology on PDA, highlighting medium-dependent phenotypes.

Carbon catabolism fuels infection and is tightly linked to virulence^[66]^. *VdGAL4* mutants show impaired use of raffinose and sucrose^[3]^; *VdOGDH* mutants grow poorly on xylan, sucrose, starch or galactose^[67]^, whereas *ΔVdHP1* over-utilises cellulose and starch^[68]^. Here, *ΔVdCut1* grew poorly on sucrose but faster on galactose, and *ΔVdPL16* was severely inhibited on sucrose or cellulose. Thus, *VdCut1* and *VdPL16* modulate carbon use and hence vegetative growth.

Fungi possess cell-wall integrity (CWI) pathways that counter abiotic stresses^[69]^. Some genes enhance both development and stress tolerance^[35,70]^. Deletion of *VdSte11* or *VdSsk2* increases sensitivity to NaCl and sorbitol^[47]^; *VdUGP* loss renders cells SDS and NaCl-hypersensitive^[71]^. Our mutants were hypersensitive to sorbitol, CR and SDS, indicating that *VdCut1* and *VdPL16* coordinate stress responses.

Penetration of cellophane membranes is an established proxy for plant root penetration^[72,73]^. *VdNoxB*, *VdPls1* and *VdSte11* mutants that fail to penetrate membranes are avirulent^[47,74]^. *ΔVdCut1* completely lost penetration ability, whereas *ΔVdPL16* retained partial but reduced activity. Correspondingly, both mutants were attenuated in cotton, with *ΔVdCut1* being more severely affected, consistent with their roles as virulence factors. Secreted effectors modulate host immunity^[75]^. Yeast signal-trap assays confirmed that *VdCut1* and *VdPL16* are secreted. Carbohydrate-active enzymes can act as PAMPs or effectors; *V. dahliae* GH12 members *VdEG1* and *VdEG3* differentially trigger immunity^[76]^. Whether *VdCut1* and *VdPL16* function as bona-fide effectors remains to be tested, but their secretion and enzymatic activity make this a promising avenue.

In summary, our results indicate that the cell wall-degrading enzyme genes *VdCut1* and *VdPL16* in *V. dahliae* regulate mycelial vegetative growth, conidial production, and carbon source utilization. They also play a role in regulating responses to abiotic stress. Furthermore, knocking out *VdCut1* and *VdPL16* genes affects mycelial penetration ability on cellophane, with a more pronounced decrease in penetration ability observed in the *VdCut1* deletion mutant. The *VdCut1* and *VdPL16* genes have a certain impact on the infection and colonization of *V. dahliae*. Overall, *VdCut1* and *VdPL16* genes play crucial roles in the infection and pathogenicity processes of *V. dahliae*.

## Acknowledgments

We are grateful to Hui-Shan Guo from the Institute of Microbiology, Chinese Academy of Sciences, for the generous gift of the vectors for gene complementation.

## Data Availability Statement

The data presented in this study are available within the article and the supplementary materials.

## Author contributions

**Conceptualization:** Gaijie Liu,Jianwei Cao.

**Data curation:** Gaijie Liu, Jingyi Ye.

**Formal analysis:** Gaijie Liu, Yulong Zhang, Xingxing Liu.

**Funding acquisition:** Aiying Wang, Asigul Ismayil.

**Investigation:** Yulong Zhang, Jianwei Cao, Jingyi Ye.

**Methodology:** Gaijie Liu, Jianwei Cao, Xingxing Liu,Yulong Zhang

. **Project administration:** Gaijie Liu, Asigul Ismayil, Aiying Wang.

**Resources:**Gaijie Liu, Asigul Ismayil.

**Supervision:** Asigul Ismayil, Aiying Wang.

**Validation:** Gaijie Liu, Aiying Wang.

**Writing – original draft:** Gaijie Liu, Jianwei Cao.

**Writing – review & editing:** Gaijie Liu, Xingxing Liu, Yulong Zhang, Asigul Ismayil, Aiying Wang.

